# Soil variation among natural habitats alters glucosinolate content in a wild perennial mustard

**DOI:** 10.1101/2022.08.16.504215

**Authors:** Maggie R. Wagner, Thomas Mitchell-Olds

## Abstract

Baseline levels of glucosinolates—important defensive phytochemicals in Brassicaceous plants—are determined by both genotype and the environment. However, the ecological causes of glucosinolate plasticity are not well characterized. Fertilization is known to alter glucosinolate content of *Brassica* crops, but the effect of naturally-occurring soil variation on glucosinolate content of wild plants is unknown. Here, we conduct greenhouse experiments using *Boechera stricta* to ask 1) whether soil variation among natural habitats shapes leaf and root glucosinolate profiles; 2) whether such changes are caused by abiotic soil properties, soil microbes, or both; and 3) whether soil-induced glucosinolate plasticity is genetically variable.

Total glucosinolate quantity differed up to two-fold between soils from different natural habitats, while the relative amounts of different compounds was less responsive. This effect was due to physico-chemical soil properties rather than microbial communities. We detected modest genetic variation for glucosinolate plasticity in response to soil. In addition, glucosinolate composition, but not quantity, of field-grown plants could be accurately predicted from measurements from greenhouse-grown plants. In summary, soil alone is sufficient to cause plasticity of baseline glucosinolate levels in natural plant populations, which may have implications for the evolution of this important trait across complex landscapes.

## Introduction

Soil properties—both abiotic and biotic—are crucial and inescapable parts of a plant’s habitat. The texture, chemical composition, and organic matter content of the local soil influence the availability of water, nutrients, and toxins to the plant, which simultaneously interacts with a highly complex community of bacteria, fungi, arthropods, nematodes, and other soil-dwelling organisms. Together, these elements affect plants’ health, appearance, and reproductive success. They also can vary among natural habitats, raising the possibility that soil characteristics influence the expression and evolution of plant phenotypes. Several examples of local adaptation to specific soil properties are well-documented—for instance, heavy metal tolerance and salinity tolerance (Macnair, 1987; Flowers and Colmer, 2008). However, due to its importance for whole-plant health and nutrition, soil quality could potentially affect the expression and evolution of a wide variety of plant traits, including in aboveground organs.

Here, we investigate the role of naturally occurring biotic and abiotic soil variation on the accumulation and evolution of glucosinolates, an important class of defensive phytochemicals produced by the Brassicales. Glucosinolates are nitrogen- and sulfur-rich compounds derived from amino acids via well-characterized biosynthetic pathways (Gigolashvili *et al*., 2009). These compounds protect against natural enemies (Mauricio and Rausher, 1997; Benderoth *et al*., 2006; Prasad *et al*., 2012) and also play a role in drought tolerance (Carley *et al*., 2021). Glucosinolate biosynthesis is often induced upon insect and pathogen attack; however, glucosinolates are also produced constitutively and stored in cell vacuoles, to be released and activated in case of cell disruption (Textor and Gershenzon 2009). The constitutive levels of glucosinolate accumulation are complex traits controlled by both genotype and environment (Wagner and Mitchell-Olds, 2018a), although the specific environmental stimuli shaping baseline glucosinolate profiles have not been fully characterized. Here, we focus on the relationship between soil and glucosinolate plasticity in natural settings. A great deal of research has explored how various agricultural soil management strategies affect glucosinolate content of *Brassica* crops, but little is known about how soils in natural habitats affect the glucosinolate profiles of wild plants.

In agricultural settings, abiotic soil variation alters glucosinolate content of both leaves and roots. Because these compounds are rich in nitrogen and sulfur, glucosinolate profiles are particularly sensitive to the availability of these two nutrients (Aires *et al*., 2006). For instance, sulfur fertilization generally increases leaf glucosinolate content, in some cases by as much as 20- to 50-fold (Falk *et al*., 2007). In addition to nitrogen and sulfur, the availability of potassium and various micronutrients can also affect glucosinolate accumulation (Shelp *et al*., 1993; Troufflard *et al*., 2010; Kim and Juvik, 2011), as can abiotic stresses such as drought, heat, and high salinity (López-Berenguer *et al*., 2008; Schreiner *et al*., 2009; Cocetta *et al*., 2018). Different glucosinolate compounds often respond differently to these abiotic stimuli; furthermore, these effects can vary among plant cultivars, developmental stages, and organs (Aires *et al*., 2006). The impact of one nutrient or soil property on glucosinolate content may also depend on the status of other nutrients or soil properties (Omirou *et al*., 2009). The complex interactions between these factors suggest that experimental transplants of entire soils can complement experimental manipulations of individual soil properties. Together, these holistic and reductionist approaches can reveal how the soils found in natural habitats affect glucosinolate accumulation in wild plants.

In addition to these abiotic components of soil variation, the microbial communities inhabiting soils from different environments vary in composition and potential functionality (Fierer, 2017). These biogeographic patterns correspond to patterns in abiotic soil properties, indicating microbial preferences for different habitats, effects of microbes on local nutrient cycling and other processes, or (most likely) some combination of these. Subsets of the soil microbiome become enriched in the rhizosphere and the interior of the root, interacting with plants via diverse biochemical mechanisms with implications for plant health and phenotype (Berendsen *et al*., 2012; Zhalnina *et al*., 2018). Pathogen attack can induce glucosinolate biosynthesis (Stahl *et al*., 2016), and the antimicrobial effects of many glucosinolate breakdown products are well documented (Kirkegaard and Sarwar, 1998; Bressan *et al*., 2009; Poveda *et al*., 2020). However, soil microbiomes also contain neutral and beneficial microbes, which can decrease or increase aliphatic glucosinolate accumulation in *Arabidopsis thaliana* depending on plant age and nutrient status (Brock *et al*., 2013; Pangesti *et al*., 2016). Furthermore, perception of microbe-associated molecular patterns in the roots stimulates expression of several genes in the glucosinolate biosynthetic pathway (Zhou *et al*., 2019). These experiments demonstrate that individual microbial strains (both pathogenic and non-pathogenic) can cause glucosinolate plasticity; however, it is unclear how such effects scale up to the level of complex soil microbiomes that vary among natural habitats.

Here, we ask 1) whether soil variation among natural habitats affects the composition and quantity of glucosinolates in a wild plant species; 2) whether such changes to glucosinolate profiles are caused by abiotic soil factors, soil microbes, or both; 3) whether plant genetic variation affects glucosinolate plasticity in response to soil; 4) how well glucosinolate profiles of greenhouse-grown plants represent those of field-grown plants of the same genotype; and 5) whether glucosinolate content interacts with either abiotic or biotic soil properties to influence seed production. To answer these questions, we conducted a series of greenhouse experiments using the wild perennial mustard *Boechera stricta* (Graham) Al-Shehbaz. Endogenous populations of *B. stricta* thrive in high-elevation forests, meadows, and riparian sites throughout the species’ native range in the Rocky Mountains (Rushworth *et al*., 2011). Because *B. stricta* is undomesticated and many of its natural habitats remain largely undisturbed, our observations of this plant’s interactions with its local soils reflect the ecological relationships that shaped its evolutionary history. Previous field studies demonstrated that *B. stricta* leaf and root glucosinolate profiles are strongly plastic both among and within habitats, and also are under natural selection (Prasad *et al*., 2012; Wagner *et al*., 2016; Wagner and Mitchell-Olds, 2018a; Carley *et al*., 2021). However, because a large number of biotic and abiotic features distinguished these habitats from each other, the ecological causes of glucosinolate plasticity and selection have not been identified in the field. In particular, little is known about how soil properties impact glucosinolate production in nature. One previous study found that soil microbial communities can protect against herbivory by inducing changes in the *A. thaliana* leaf metabolome, but did not quantify glucosinolates in particular (Badri *et al*., 2013). Another found that water deprivation altered *Boechera* glucosinolate quantity and glucosinolate profile composition, but did not investigate effects of soil *per se* (Haugen *et al*., 2008). A third reported correlations between *Brassica rupestris* glucosinolate content and soil properties including organic matter and microbe-derived carbon, but did not verify this relationship experimentally (Muscolo *et al*., 2019). Here, our results confirm that physicochemical properties of diverse soils from natural *B. stricta* habitats can have a direct effect on glucosinolate accumulation.

## METHODS

We conducted two sequential greenhouse experiments to investigate how interactions between plant genotype, soil chemistry, and soil microbes affect glucosinolate content in *B. stricta*. For the first (“whole soil”) experiment, we grew twenty *B. stricta* inbred lines in soils collected from four natural habitats in its native range, plus a potting-soil control. We then measured glucosinolate content of roots and rosette leaves of six-week-old plants.

For the second experiment, we separated the biotic and abiotic components of naturally-occurring soils and assessed their relative effects on aboveground traits. We grew a panel of 48 *B. stricta* inbred lines either in a common sterilized potting soil that had been inoculated with microbes from one of four wild soils (“biotic” experiment) or in a sterilized wild soil (“abiotic” experiment). We then measured leaf glucosinolate content, growth rate, and fruit production of adult plants.

### Origins of soils and plant genotypes

All soils used in these experiments were collected from remote locations in central Idaho, USA, where wild populations of *B. stricta* are endemic. For the first (“whole-soil”) experiment, we collected soil in August 2011 from four field sites: Mahogany Valley (“Mah”), Mill Campground (“Mil”), Parker Meadow (“Par”), and Silver Creek (“Sil”). These sites range in elevation from 1812 m to 2676 m and vary in many environmental attributes including water availability, soil chemistry, and the composition and diversity of both plant and microbial communities (Figure 1 (Wagner *et al*., 2014; Wagner and Mitchell-Olds, 2018a)). At each site, five evenly spaced subsamples (∼12 m apart, from 10 cm to 30 cm deep) were mixed, pooled, and passed through a ∼1.25 cm sieve to create the final soil sample, which was shipped to Duke University and stored at 4°C until use. For the second (“biotic/abiotic”) experiment, we collected soils in August 2012 using the same method. We sampled Mahogany Valley, Parker Meadow, and Silver Creek for this experiment as well, but for logistical reasons, we substituted a fifth field site (Jackass Meadow, or “Jam”) for Mill Campground. We also collected twelve subsamples per location (spaced approximately 1 m apart) for nutrient analysis (Figure 2c).

**Figure 1:**
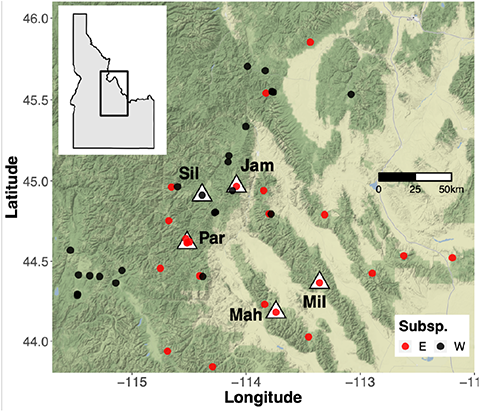
Origins of field soils and wild *B. stricta* genotypes used in two greenhouse experiments. Natural habitats from which soils were sampled are labeled and denoted with white triangles. Circles mark collection sites of the genotypes included in the experiment. Not shown: one EAST subspecies genotype collected in Colorado. Map tiles by Stamen Design, under CC BY 3.0.

**Figure 2:**
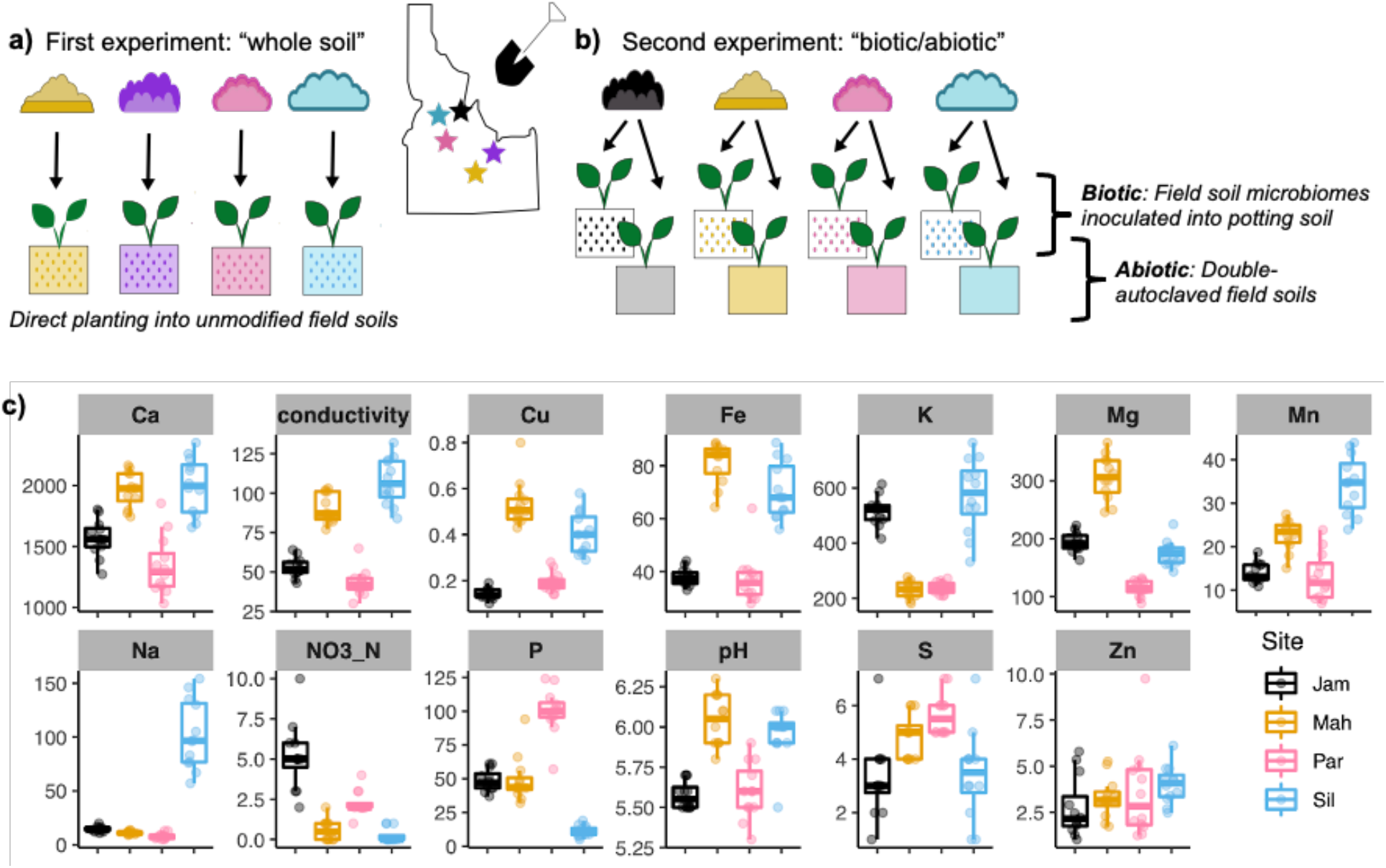
Soils from five natural *B. stricta* habitats were used for two independent greenhouse experiments. Map is not to scale. (a) In the “whole soil” experiment, individuals were planted into unmodified soils so that the experimental treatments encompassed both variation in microbial communities (symbolized by dots) and in physico-chemical soil properties (symbolized by solid color backgrounds). (b) In the second experiment, field soils were separated to create separate sets of four “biotic” treatments (soil microbiomes from each habitat in a potting soil matrix) and four “abiotic” treatments (sterilized soil matrices from each habitat). (c) Nutrient analysis of field soils used in the second experiment. N=12 subsamples from each field site, approximately 1 m between subsamples. pH is unitless; units for conductivity are µmho/cm; all other variables have units of ppm. Data were generated by the Texas A&M AgriLife Extension Service Soil, Water, and Forage Testing Laboratory.

All *B. stricta* genotypes used in both experiments were inbred lines descended from accessions collected from wild populations in the species’ native range (Figure 1). These lines were propagated via self-pollination in the greenhouse for at least one generation to reduce maternal effects. Because *B. stricta* is naturally self-fertilizing and highly inbred (Song *et al*., 2006), the seeds collected from a single mother plant are self-full siblings that are essentially genetic clones. Therefore, the use of these lines allowed us to deduce that any phenotypic variation among individuals of the same genotype was caused by environmental variation, i.e., phenotypic plasticity in response to soil treatment.

For the first experiment, we selected a subset of the lines that had been used in a parallel field experiment (Wagner *et al*., 2016). This group comprised twenty lines total; four from each of five wild populations endemic to the five field sites from which we collected soil (Figure 1). Because *B. stricta* is characterized by high *F*_ST_ (i.e., high genetic divergence among populations but low levels of genetic variation within populations), the four lines within each population were more genetically similar to each other than they were to the lines from the other populations (Song *et al*., 2006). Because of this, and for consistency with the second experiment (described below) and with previous experiments (Wagner *et al*., 2016; Wagner and Mitchell-Olds, 2018a), we define “genotype” as any group of accessions collected from the same wild population, as well as the lines descended from these accessions via self-pollination.

For the second experiment, we used a more diverse panel of 48 *B. stricta* genotypes, each of which was derived via single-seed descent from one accession collected from one wild population (Wagner *et al*., 2014). Unlike the group of five genotypes used in the first experiment, these 48 genotypes were evenly split between the EAST and WEST subspecies, which are genetically and ecologically divergent but occupy partially overlapping parts of the species range (Figure 1; (Lee and Mitchell-Olds, 2013)). One line per genotype was used for this experiment.

### Soil treatments

For the first (“whole-soil”) experiment (Figure 2a), we used a split-plot design by filling three 48-cell trays with each field-collected soil. Each pot in the tray measured 1.5 in wide, 2.5 in long, and 2.25 in deep. We filled another three trays with potting soil: Fafard 4P (Conrad Fafard Inc., Agawam, MA, USA) on the bottom 75%, and Metromix 200 (Sun Gro Horticulture Inc., Vancouver, BC, Canada) on the top 25%.

For the second experiment, we modified field-collected soils to separate them into their “biotic” and “abiotic” components (Figure 2b). To create the biotic soil treatments, we stirred 75 g of each field soil into 1 L of sterile MES monohydrate (2.5 mM; Sigma Aldrich, St. Louis, MO, USA). We allowed the suspensions to settle for 30 minutes and then vacuum-filtered them through sterile filter paper with 11-µm pores. We then centrifuged the filtrates at 3,000 x *g* for 30 minutes. We discarded the supernatants and resuspended the pellets—which contained soil microorganisms—in 1L of sterile 2.5 mM MES monohydrate. We diluted 400 mL of each microbial slurry along with 6 g of 20-10-20 fertilizer into 4 L of sterile diH_2_O, which we then used to saturate 49-mL Conetainer pots (Stuewe & Sons, Inc., Tangent, OR, USA) filled with double-autoclaved potting soil (121°C for 90 minutes, separated by 24 hours at room temperature). The potting soil included 75% Fafard 4P on the bottom layer, and a top layer of Metromix 200. We added an additional 1 mL of undiluted slurry to the top of each pot via pipette. In total, each of the 4 microbial treatments was applied to 200 pots. Because the same potting-soil medium was used to create all four “biotic” treatments, these soils differed in microbial content but were otherwise identical. We cannot rule out that the filtering process and subsequent recolonization resulted in soil microbiomes that are somewhat different from those found in the original wild soils. However, the differences between our four inocula still reflect the profound variation in soil community composition among natural *B. stricta* habitats (Wagner *et al*., 2014, 2016).

To create the “abiotic” soil treatments, we double-autoclaved the remainder of the field-collected soils (121°C for 90 minutes, separated by 24 hours at room temperature). After mixing in ¼ volume of autoclaved perlite to improve drainage, we filled 200 Conetainer pots with each of the four sterilized field soils. Like for the biotic soil treatments, individual pots of the different treatments were fully randomized among racks before planting.

### Plant care and phenotyping

For both experiments, we surface-sterilized seeds by vortexing them in 70% ethanol with 0.1% Triton X-100 for 1 minute, vortexing and incubating them in 10% bleach with 0.1% Triton X-100 for 15 minutes, and then rinsing them three times in sterile diH_2_O. We placed surface-sterilized seeds onto autoclaved filter paper with sterile diH_2_O in sterile petri dishes sealed with Parafilm. The seeds were stratified for one week in the dark at 4°C and then transferred to a growth chamber (22°C, 11-hour days, ambient humidity). After allowing seeds to germinate for one week, we moved them to the greenhouse and transplanted them into their assigned soil treatments.

For the first experiment, we planted seven replicates per line (i.e., 28 replicates per genotype) in each soil type, randomized across the three blocks per soil type. Four unplanted pots per soil type were kept as controls. Plants received tap water as needed, and the placement of trays within the greenhouse was randomized and regularly rotated. Plants grew under standard greenhouse conditions for the duration of the experiment (daytime temperature 65-70°F with 600–2,000 µmol s^-1^ cm^-2^ photosynthetically active radiation, nighttime temperature 55-60°F, relative humidity maintained between 37 and 52%). When plants were six weeks old, we harvested ∼30 mg each of leaf tissue and root tissue into tubes of 70% methanol for glucosinolate extraction.

For the second experiment, we planted four individuals per genotype into each of the eight soil treatments, arranged in fully randomized blocks of 200 pots. Eight pots per treatment were left unplanted. We top-watered plants with reverse osmosis (RO) water as needed for the duration of the experiment. When the plants were one month old, we applied an additional 4 mL of 20-10-20 fertilizer in sterile diH_2_O to each pot via pipet. Plants grew under standard greenhouse conditions except for a seven-week vernalization treatment (4°C) when the plants were 2 months old. For eight weeks after the end of vernalization, we checked each plant’s reproductive status three times per week. When an individual’s first open flower was observed, we measured plant size (height and number of leaves) and collected ∼30 mg of rosette leaf tissue into 70% methanol for glucosinolate extraction (see below). For the 749 plants that did not flower within eight weeks after vernalization, we made these measurements and collected leaf tissue on the day of the last flowering census. Plants were allowed to mature for eight weeks after the last flowering census, at which point we measured fecundity by counting the number of fruits produced by each individual. Because destructive sampling of roots would have prevented later measurement of fruit output, and because our primary goal was to understand the ecological causes of leaf glucosinolate plasticity among habitats (Wagner and Mitchell-Olds, 2018a), we did not measure root glucosinolates for this experiment.

### Measurement of glucosinolate content

We extracted glucosinolates by passing the 70% methanol leachates through columns of DEAE-Sephadex A-25 (Sigma Aldrich, St. Louis, USA) that had been equilibrated in 20 mM sodium acetate for one hour. We added 50 *µ*L of 1 mM sinigrin (Sigma Aldrich, St. Louis, USA) to each column as an internal standard. We randomized all samples across 96-well plates, each of which also contained methanol and sinigrin-only negative controls. The columns were washed repeatedly (2 x 750 *µ*L 70% methanol; 2 x 750 *µ*L diH_2_O; 1 x 750 *µ*L 20mM sodium acetate) and then incubated overnight in sulfatase. Desulfinated glucosinolates were then eluted in four 75 *µ*L fractions: two in 70% HPLC-grade methanol, and two in HPLC-grade water. These fractions were pooled prior to analysis. We analyzed 50 *µ*L of each sample using an Agilent 1100-series HPLC machine with a diode array detector and a Zorbax Eclipse XDB-C18 column (4.6 x 150 mm, pore size 5 *µ*m; Agilent Technologies, Santa Clara, USA). Desulfinated glucosinolates were separated at 40°C in a mobile phase beginning with 1.5% acetonitrile (ACN) in HPLC-grade water for 6 minutes and then increasing linearly to 2.5% ACN over two minutes; to 5% ACN over 7 minutes; to 18% ACN over 2 minutes; to 46% ACN over 6 minutes; to 92% ACN over 1 minute; and finally decreasing to 1.5% ACN over 5 minutes. Individual compounds were identified using their retention times and their UV absorption spectra at 229 nm (Windsor *et al*., 2005; Olson-Manning *et al*., 2013).

We calculated the absolute concentration (µmol/mg) of each compound in each sample as:

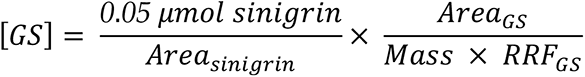

*Area_sinigrin_* denotes the area under the sinigrin chromatograph peak; *Area_GS_* denotes the area under the peak corresponding to the focal compound; *Mass* refers to the air-dried mass (measured in mg) of the tissue sample used for glucosinolate extraction; and *RRF_GS_* denotes the relative response factor of the focal compound (Brown *et al*., 2003; Clarke, 2010).

In the “whole-soil” experiment, most samples contained some combination of the indolic compound 4-methoxyindol-3-ylmethylglucosinolate (4MOI3M) and five aliphatic compounds: 2-hydroxy-1-methylethylglucosinolate (2OH1ME), 1-methylethylglucosinolate (1ME), 6-methylsulfinylhexylglucosinolate (6MSOH), 1-methylpropylglucosinolate (1MP), and 6-methylthiohexylglucosinolate (6MTH; Figure 3). The large number of samples in the “biotic/abiotic” experiment (N>1000) made it infeasible to measure 6MTH and 4MOI3M, which due to their low polarity required additional time and reagents to quantify (see below, “Identification of two late-eluting glucosinolates”). However, because these two compounds were only minor components of leaf glucosinolate profiles (Figure 3), and because the “biotic/abiotic” experiment only investigated leaves, their exclusion from this dataset should not have a major impact on our conclusions.

**Figure 3:**
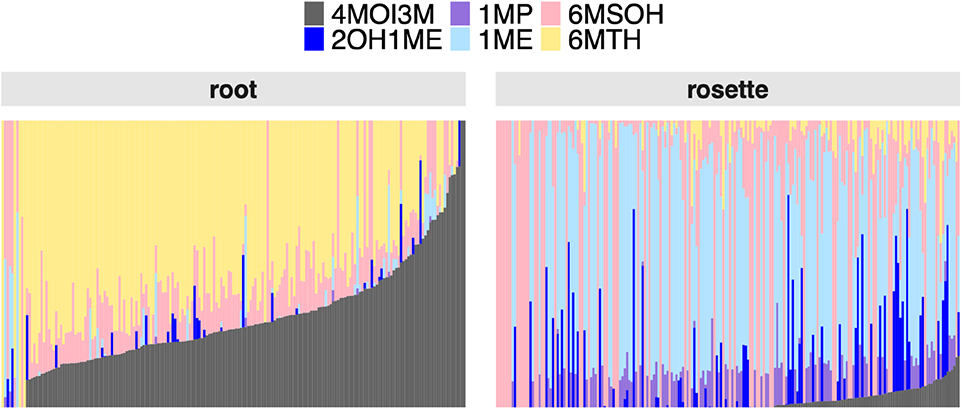
Glucosinolate profiles of 403 samples from greenhouse-grown plants in natural soils (*i.e.,* the “whole-soil” experiment). Each vertical bar shows the proportional abundance of the six compounds measured in this experiment. 6MSOH and 6MTH are methionine-derived aliphatic glucosinolates; 2OH1ME, 1ME, and 1MP are branched-chain aliphatic glucosinolates; and 4MOI3M is an indolic glucosinolate.

We used the absolute concentrations of these six compounds to calculate three summary metrics that describe emergent properties of glucosinolate profiles:

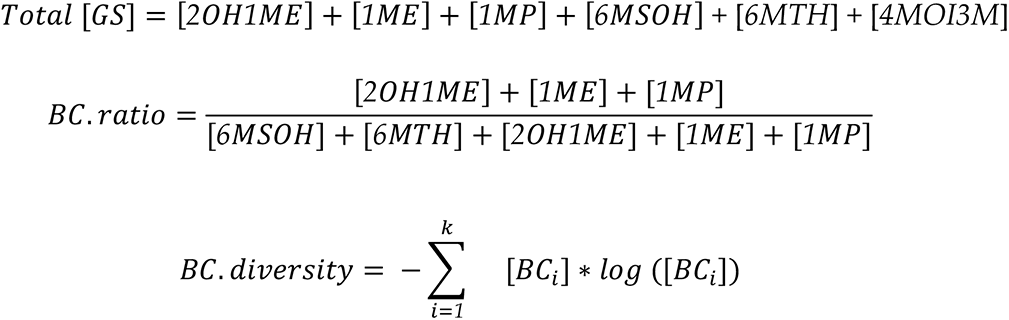

Total [GS] is the summed abundance of all glucosinolate compounds. BC-ratio, the proportion of aliphatic compounds with branched-chain structures, is key to insect resistance and is evolving non-neutrally in *B. stricta* (Schranz *et al*., 2009; Manzaneda *et al*., 2010; Prasad *et al*., 2012; Carley *et al*., 2021). Finally, BC-diversity is the Shannon diversity index applied to the three branched-chain aliphatic glucosinolates. In the formula, *k* is the number of branched-chain glucosinolate types and [*BC_i_*] is the absolute concentration of the *i^th^* branched-chain compound. Therefore, BC-diversity describes the “evenness” of these compounds: low values indicate that one compound dominates a glucosinolate profile and high values indicate that multiple compounds are equally abundant. Previous work has shown that all of these emergent properties of glucosinolate profiles show high phenotypic plasticity both within and among natural habitats, and are associated with reproductive fitness in nature (Wagner and Mitchell-Olds, 2018a).

### Identification of two late-eluting glucosinolates

We used leftover desulfoglucosinolate mixtures from a subset of the samples to identify two late-eluting compounds that were commonly observed in the “whole-soil” experiment but had not been previously identified in *B. stricta*. High-pressure liquid chromatography with diode-array detector was performed as described earlier (Burow *et al*., 2006) using Agilent 1100 high-pressure liquid chromatography with 96-well plate autosampler. The eluted desulfoglucosinolates were separated on a reversed phase C-18 column (Nucleodur Sphinx RP, 250 x 4.6 mm, 5µm, Macherey-Nagel, Düren, Germany) with a water (A)-acetonitrile (B) gradient (0-1 min, 1.5% B; 1-6 min, 1.5-5% B; 6-8 min, 5-7% B; 8-18 min, 7-21% B; 18-23 min, 21-29% B; 23-23.1 min, 29-100% B; 23.1-24min 100% B and 24.1-28 min 1.5% B; flow 1.0 mL min-1). Identity of desulfoglucosinolates was confirmed by liquid chromatography-electrospray mass spectrometry on a Bruker Esquire 6000 ion trap mass spectrometer (Bruker Daltonics, Bremen, Germany) with the same chromatographic conditions.

We identified 4-methoxyindol-3-ylmethylglucosinolate (4MOI3M) based on match of retention time of the desulfoglucosinolate in the sample with that of known desulfo-4-methoxyindol-3-ylmethylglucosinolate from a leaf extract of *Arabidopsis thaliana* Col-0. The full scan mass spectrum in positive mode (base peak m/z 399; insource fragment m/z 237) and the UV spectrum in the range 190 to 360 nm also matched.

We also identified 6-methylthiohexylglucosinolate (6MTH) on the basis of its desulfo-derivative. The full scan mass spectrum in positive mode (base peak m/z 370; insource fragment m/z 208) matched desulfo-6-methylthiohexylglucosinolate and the UV spectrum in the range 190 to 360 nm matched that of desulfo-methylthioalkylglucosinolate.

### Statistical analysis

For the first (“whole-soil”) experiment, low survival precluded robust analysis of the MIL genotype and the potting-soil treatment; therefore, we excluded these groups from statistical analyses. After this filtering, we performed separate analyses of glucosinolate data for roots (*N*=161) and rosettes (*N*=172). We modeled each glucosinolate trait (Total [GS], BC-ratio, and BC-diversity) using a REML mixed-effects model:

*Trait* = *Genotype* + *Soil* + *Genotype* ∗ *Soil* + *Line*[*Genotype*] + *Block*[*Soil*] + *Batc*ℎ where *Line* (nested in *Genotype*), *Block* (nested in *Soil*), and *Batch* were random-intercept terms. In cases where the full model resulted in a singular fit, we removed the least informative random-intercept term(s) to fix the singularity.

For the second experiment, we separately analyzed the “biotic” (*N*=476) and “abiotic” (*N*=539) treatment sets. We modeled each trait (Total [GS], BC-ratio, and BC-diversity) using a REML mixed-effects model:

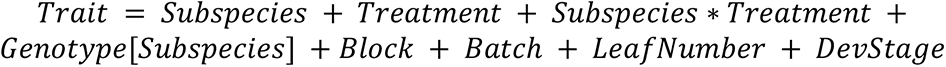

where *Genotype* (nested in *Subspecies*), *Block*, and *Batch* were random-intercept terms. *Leaf Number* and *Developmental Stage* were covariates to controlfor effects of overall plant size and ontogenetic status on rosette glucosinolate content (Agrawal, 2011). As above, in cases where the full model resulted in a singular fit, we removed the least informative random-intercept term(s) to fix the singularity.

Finally, we used genotypic selection analysis (Rausher, 1992) to ask whether patterns of selection on glucosinolate traits differed among soil treatments. The dependent variable for these models was the relative fecundity of each genotype in each treatment (*genotype’s mean fecundity / grand mean fecundity of all genotypes*). The independent variables were the mean trait values of each genotype in each treatment. To test for directional (linear) selection on each trait, we used the model:

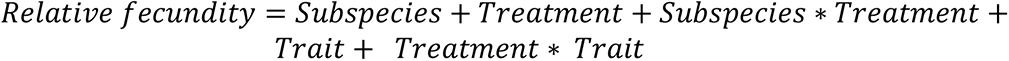

where the linear coefficient of the *Trait* term describes the average selection differential across all treatments, and the *Treatment * Trait* interaction indicates whether selection varied among treatments. To test for stabilizing or disruptive (quadratic) selection on each trait, we fit separate models with additional quadratic terms: *Trait*^2^ + *Treatment * Trait^2^*. We assessed the significance of all terms using ANOVA with Type III sums of squares.

For both experiments, *Batch* was a nuisance variable to account for differences in efficiency of glucosinolate extractions or HPLC measurements among randomized plates of samples. When used as dependent variables, Total [GS] was square-root transformed and BC-ratio was arcsine-square-root transformed to improve homoscedasticity. We used likelihood ratio tests to determine statistical significance of random effects, and Type III ANOVA to assess fixed effects, with Satterthwaite approximation of degrees-of-freedom. We used **R** version 4.2.0 for all analyses and figures, particularly the packages tidyverse, lme4, lmerTest, and emmeans (Bates *et al*., 2015; Kuznetsova *et al*., 2017; Wickham *et al*., 2019; Lenth 2022). Whenever appropriate, we used the sequential Bonferroni correction to adjust *P-*values for multiple comparisons.

### Comparison to data from field experiments

To compare glucosinolate content between greenhouse-grown and field-grown plants, we combined our data from the experiments described above with two published datasets from field experiments that used the same genotypes (Wagner *et al*., 2017 for the five whole-soil experiment genotypes; Wagner and Mitchell-Olds, 2018b for the biotic/abiotic experiment genotypes). We subsetted each dataset to include only the genotypes and habitats represented in both experiments. Interpretation of these comparisons is not straightforward because the field-grown plants were up to 3 years older than the greenhouse-grown plants, and because the field-grown plants were started in the greenhouse and then transplanted into the field sites in plugs of potting soil. Despite these complications, it is useful to assess how well greenhouse measurements predict trait values in the field.

Due to the small number of genotypes in the whole-soil experiment, we relied on qualitative comparisons of both root and shoot data. For the biotic/abiotic experiment, however, we calculated genetic correlations between field-based and greenhouse-based measurements of each trait. To do this, we fit fixed-effects linear models:

*Trait* = *Genotype* + *Experiment* + *Genotype* ∗ *Experiment* where *Experiment* was either “biotic”, “abiotic”, or “field”. Estimated marginal means for each genotype in each experiment were derived from this model. Then, we regressed the genotype means from field data onto the genotype means from greenhouse data, separately for the biotic and abiotic experiments.

Finally, for a holistic assessment of the major drivers of leaf glucosinolate variation, we used the **vegan** package (Oksanen *et al*. 2020) to conduct an unconstrained principal components analysis (PCA) of the four aliphatic compounds measured in the biotic, abiotic, and field experiments. Technical noise due to Batch and Block effects was partialled out prior to the unconstrained ordination.

## RESULTS

### Natural soil variation is sufficient to alter glucosinolate quantity, but not quality

For the “whole-soil” greenhouse experiment, we measured six different glucosinolates in both leaves and roots of plants grown in four different field-collected soils. In general, glucosinolate profiles were dominated by aliphatic compounds in leaves, but by indolic compounds in roots—the same pattern that has been observed in *Brassica* crops (Figure 3; (Rosa, 1997; Aires *et al*., 2006; Omirou *et al*., 2009)). The total concentration of aliphatic glucosinolates was positively correlated with concentration of the indole 4MOI3M in both leaves (*r*=0.27, *P*=6.8e-5) and roots (*r*=0.58, *P*=1.5e-18), justifying the pooling of all six compounds into a single emergent trait, Total [GS]. Within individual plants, there was no correlation between Total [GS] in leaves and roots (*r*=0.021, *P*=0.80), suggesting that glucosinolate production is regulated independently in these organs. In contrast, both BC-ratio (*r*=0.35, *P*=7.0e-6) and BC-diversity (*r*=0.19, *P*=0.017) were weakly but positively correlated between roots and leaves.

Consistent with previous studies of *B. stricta*, variation among genotypes and among lines within genotypes strongly influenced both BC-ratio and BC-diversity in leaves (Table 1; Figure 4b; Schranz *et al*., 2009; Wagner and Mitchell-Olds, 2018a. However, the lack of variation among lines for root glucosinolate profiles in this experiment differs from previous observations from field experiments (Wagner *et al*., 2016), indicating either that the genetic variation is expressed in natural but not greenhouse conditions, or that the relatively small subset of lines analyzed in the greenhouse experiment (16 lines as opposed to 48 in the field experiment) was not sufficient to capture this effect in roots.

**Figure 4:**
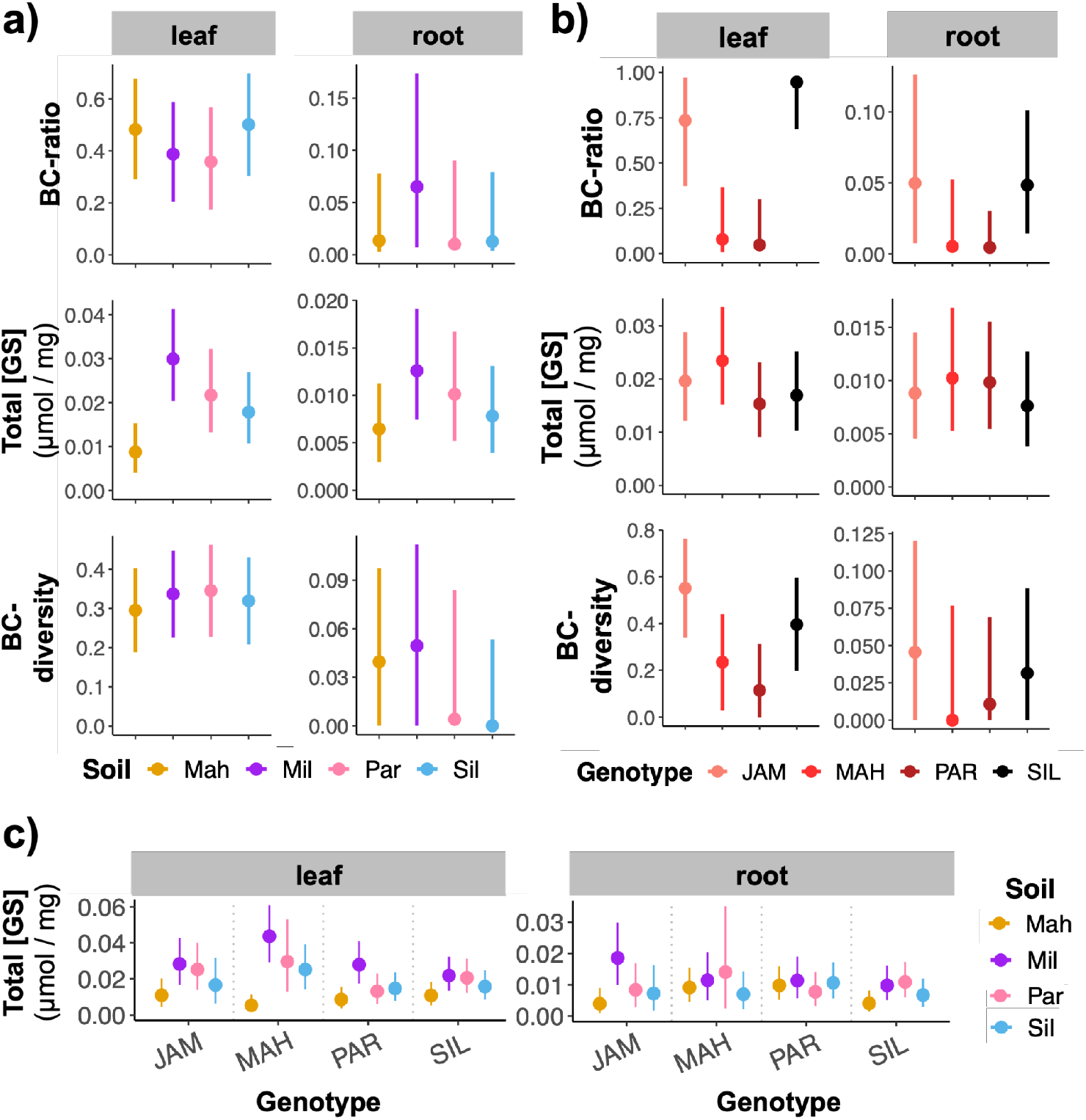
Variation in glucosinolate traits attributed to a) soil, b) genotype, and c) their interaction. Estimated marginal mean trait values with 95% confidence intervals are shown for three emergent traits of glucosinolate profiles: branched-chain ratio, branched-chain diversity, and total glucosinolate concentration. Statistics associated with these results are found in Table 1.

**Table 1:** Statistics from REML mixed models of three glucosinolate traits in leaves and roots of plants grown in whole natural soils. Values of *F* and *Χ^2^* are shown for fixed and random effects, respectively. All *P* values were corrected for multiple comparisons. Empty cells indicate that the random-intercept term was uninformative, caused a singular model fit, and was omitted from the final model.

|  |  | BC-ratio | Total [GS] | BC-diversity |
| --- | --- | --- | --- | --- |
| leaf | model $R^2$ | 0.94 | 0.57 | 0.83 |
| | Soil | $F_{3,10} = 2.26$<br>$P = 0.29$ | $F_{3,13} = 14.57$<br>$P = 0.00051$ | $F_{3,14} = 0.81$<br>$P = 0.51$ |
| | Genotype | $F_{3,13} = 11.3$<br>$P = 0.002$ | $F_{3,149} = 2.80$<br>$P = 0.061$ | $F_{3,13} = 4.09$<br>$P = 0.06$ |
| | Soil x<br>Genotype | $F_{9,139} = 1.18$<br>$P = 0.32$ | $F_{9,149} = 2.79$<br>$P = 0.014$ | $F_{9,140} = 1.54$<br>$P = 0.28$ |
| | Line<br>[Genotype] | $X^2_1 = 173.54$<br>$P = 2.5e-39$ | | $X^2_1 = 131.83$<br>$P = 1.63e-30$ |
| | Block<br>[Soil] | $X^2_1 = 2.39$<br>$P = 0.37$ | $X^2_1 = 1.02$<br>$P = 0.62$ | $X^2_1 = 0.53$<br>$P = 0.62$ |
| | Batch | $X^2_1 = 3.99$<br>$P = 0.091$ | $X^2_1 = 17.16$<br>$P = 1.03e-4$ | $X^2_1 = 0.2$<br>$P = 0.65$ |
| root | model $R^2$ | 0.34 | 0.32 | 0.30 |
| | Soil | $F_{3,9} = 0.94$<br>$P = 0.80$ | $F_{3,141} = 4.24$<br>$P = 0.020$ | $F_{3,10} = 1.09$<br>$P = 0.80$ |
| | Genotype | $F_{3,15} = 4.08$<br>$P = 0.079$ | $F_{3,141} = 1.33$<br>$P = 0.54$ | $F_{3,7} = 0.45$<br>$P = 0.73$ |
| | Soil x<br>Genotype | $F_{9,134} = 1.04$<br>$P = 0.83$ | $F_{9,141} = 1.90$<br>$P = 0.17$ | $F_{9,130} = 0.89$<br>$P = 0.83$ |
| | Line<br>[Genotype] | $X^2_1 = 0.43$<br>$P = 0.51$ | | $X^2_1 = 2.99$<br>$P = 0.17$ |
| | Block<br>[Soil] | $X^2_1 = 8.29$<br>$P = 0.008$ | | $X^2_1 = 2.5$<br>$P = 0.11$ |
| | Batch | | $X^2_1 = 15.05$<br>$P = 1.05e-4$ | |

Soil had a strong main effect on glucosinolate concentration, but not on BC-ratio or BC-diversity (Table 1). Most strikingly, the soil from Mahogany Valley caused a strong decrease in glucosinolate quantity in both leaves and roots, while the Mill Campground soil generally boosted glucosinolate quantity (Figure 4a). This result is consistent with previous observations from the field, where plants growing in Mahogany Valley had lower Total [GS] than plants growing in the other habitats (Wagner and Mitchell-Olds, 2018a). This shows that plasticity of glucosinolate quantity across a natural landscape is at least partly caused by soil variation. However, based on this experiment, the plasticity of leaf BC-ratio, leaf BC-diversity, and root BC-ratio in the field are more likely caused by other environmental factors that distinguish those habitats (Wagner *et al*., 2016; Wagner and Mitchell-Olds, 2018a).

Finally, we detected genotype-by-soil interactions driving Total [GS] in leaves, but not in roots (Table 1). The phenotypic response to soil variation was strongest in the genotype from Mahogany Valley and weakest in the genotype from Silver Creek (Figure 4c), which was the only genotype from the WEST subspecies included in this experiment. This pattern is congruent with observations from the field, where glucosinolate profiles of EAST genotypes were much more plastic across the landscape than those of WEST genotypes (Wagner and Mitchell-Olds, 2018a).

### Glucosinolate plasticity in response to soil variation is primarily driven by abiotic factors

To further dissect the ecological causes of glucosinolate plasticity in leaves of *B. stricta*, we separated four natural soils into their biotic and abiotic components. We measured aliphatic glucosinolates in leaves of 48 *B. stricta* genotypes grown in four different sterilized field soils (the “abiotic” experiment, *N* = 539) and in sterilized potting soil inoculated with microbial communities from the same four wild soils (the “biotic” experiment, *N* = 476; Figure 2). The field soils were chemically distinct: except for zinc, every soil variable that we measured differed significantly among habitats (Figure 2c; all *P*<0.001 by ANOVA after sequential Bonferroni correction).

All three glucosinolate traits (BC-ratio, BC-diversity, and Total [GS]) were highly variable among genotypes, consistent with results from field experiments (Table 2; (Wagner and Mitchell-Olds, 2018a)). On average, the WEST subspecies had higher BC-diversity and higher Total [GS] than the EAST subspecies; however, these differences were only detectable in the “abiotic” experiment (Figure 5c-d). This suggests that some genetic differences between the subspecies were not expressed in plants growing in potting soil rather than natural substrates.

**Figure 5:**
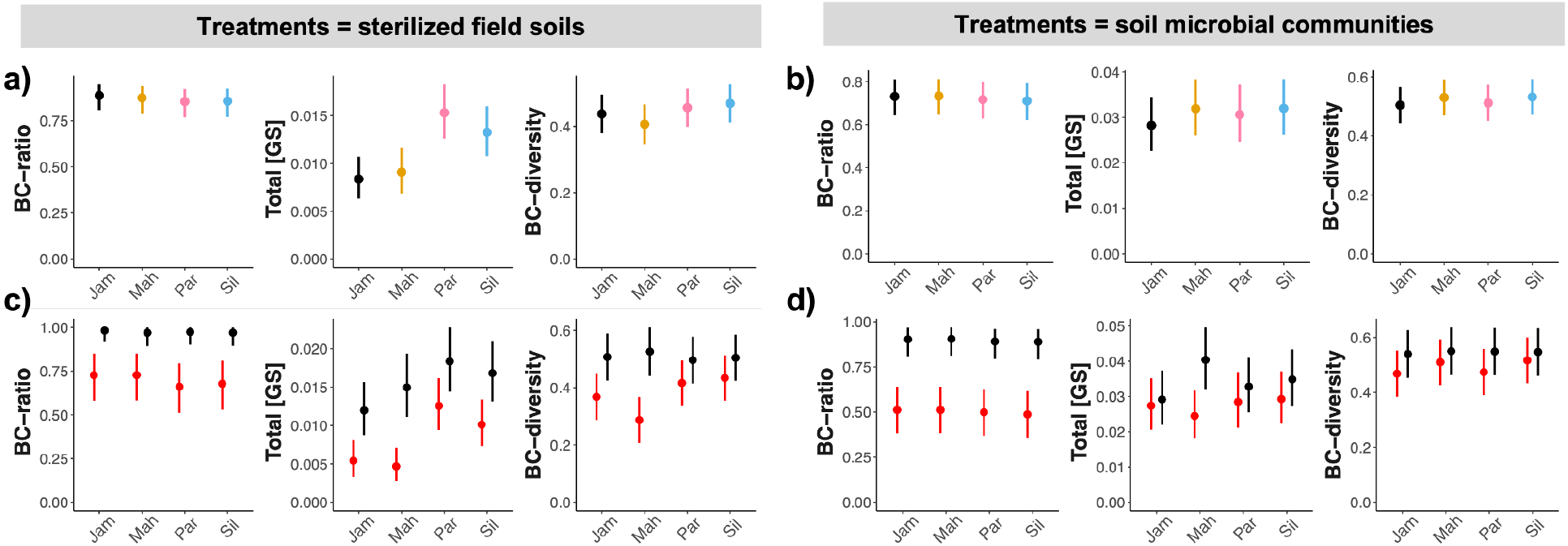
Variation in leaf glucosinolate content attributed to differences among soil treatments and plant genotype. Estimated marginal mean trait values with 95% confidence intervals are shown for three emergent traits of glucosinolate profiles: branched-chain ratio, total glucosinolate concentration, and diversity of branched-chain compounds. Units for Total [GS] are µmol/mg. Panel (a) shows trait values for plants grown in sterilized soils from four natural habitats, averaged across all genotypes. Panel (b) shows results for plants grown in sterilized potting soil inoculated with microbial communities isolated from soils of the same four natural habitats. Panels (c-d) break down the responses to the soil treatments by plant subspecies; the EAST subspecies is in red, and the WEST subspecies is in black.

**Table 2:** Statistics from REML mixed models of glucosinolate traits in leaves of plants grown in (a) sterilized potting soil inoculated with one of four natural soil microbial communities, or (b) sterilized field soil collected from one of four natural habitats. Values of *F* and *Χ^2^* are shown for fixed and random effects, respectively. All *P* values were corrected for multiple comparisons. Empty cells indicate that the random-intercept term was uninformative, caused a singular model fit, and was omitted from the final model.

| (a) |  | BC-ratio | Total [GS] | BC-diversity |
| --- | --- | --- | --- | --- |
| Treatment<br>s =<br>Sterilized<br>Field<br>Soils | model $R^2$ | 0.89 | 0.44 | 0.69 |
| | Subspecies | $F_{1,46} = 15.94$<br>$P = 0.00047$ | $F_{1,48} = 19.96$<br>$P = 0.00015$ | $F_{1,47} = 6.68$<br>$P = 0.013$ |
| | Sterilized field soil | $F_{3,151} = 2.52$<br>$P = 0.060$ | $F_{3,137} = 13.74$<br>$P = 2.1\text{e-}7$ | $F_{3,154} = 3.37$<br>$P = 0.040$ |
| | Subspecies x<br>Sterilized field soil | $F_{3,131} = 1.46$<br>$P = 0.23$ | $F_{3,118} = 2.72$<br>$P = 0.095$ | $F_{3,133} = 8.22$<br>$P = 0.00014$ |
| | Leaf number | $F_{1,462} = 1.47$<br>$P = 0.45$ | $F_{1,503} = 5.1$<br>$P = 0.073$ | $F_{1,486} = 1.12$<br>$P = 0.45$ |
| | Developmental Stage | $F_{1,474} = 5.71$<br>$P = 0.052$ | $F_{1,512} = 0.91$<br>$P = 0.34$ | $F_{1,493} = 4.49$<br>$P = 0.069$ |
| | Genotype [Subsp.] | $X^2_1 = 292.7$<br>$P = 3.9\text{e-}65$ | $X^2_1 = 37.05$<br>$P = 1.2\text{e-}9$ | $X^2_1 = 146.34$<br>$P = 2.2\text{e-}33$ |
| | Genotype x Sterilized<br>field soil | $X^2_1 = 0.46$<br>$P = 1$ | $X^2_1 = 0.03$<br>$P = 1$ | $X^2_1 = 0.79$<br>$P = 1$ |
| | Block | $X^2_1 = 9.28$<br>$P = 0.0047$ | $X^2_1 = 0.99$<br>$P = 0.32$ | |
| | Batch | | $X^2_1 = 3.52$<br>$P = 0.077$ | $X^2_1 = 4.28$<br>$P = 0.077$ |
| (b) |  | BC-ratio | Total [GS] | BC-diversity |
| Treatment<br>s = Soil<br>Microbial<br>Communi<br>ties | model $R^2$ | 0.94 | 0.40 | 0.87 |
| | Subspecies | $F_{1,46} = 25.15$<br>$P = 2.5\text{e-}5$ | $F_{1,50} = 3.64$<br>$P = 0.12$ | $F_{1,46} = 0.89$<br>$P = 0.35$ |
| | Soil microbiome | $F_{3,402} = 1.59$<br>$P = 0.38$ | $F_{3,121} = 0.94$<br>$P = 0.42$ | $F_{3,416} = 2.8$<br>$P = 0.12$ |
| | Subspecies x<br>Soil microbiome | $F_{3,417} = 0.05$<br>$P = 0.98$ | $F_{3,121} = 2.89$<br>$P = 0.11$ | $F_{3,414} = 1.87$<br>$P = 0.27$ |
| | Leaf number | $F_{1,424} = 7.07$<br>$P = 0.016$ | $F_{1,442} = 5.50$<br>$P = 0.019$ | $F_{1,426} = 9.07$<br>$P = 0.0083$ |
| | Developmental Stage | $F_{1,424} = 2.10$<br>$P = 0.45$ | $F_{1,457} = 0.14$<br>$P = 0.90$ | $F_{1,423} = 0.58$<br>$P = 0.90$ |
| | Genotype [Subsp.] | $X^2_1 = 831.9$<br>$P = 1.9\text{e-}182$ | $X^2_1 = 36.08$<br>$P = 1.9\text{e-}9$ | $X^2_1 = 671.12$<br>$P = 1.1\text{e-}147$ |
| Treatment<br>s =<br>Sterilized<br>Field<br>Soils | model $R^2$ | 0.89 | 0.44 | 0.69 |
| | Subspecies | $F_{1,46} = 15.94$<br>$P = 0.00047$ | $F_{1,48} = 19.96$<br>$P = 0.00015$ | $F_{1,47} = 6.68$<br>$P = 0.013$ |
| | Sterilized field soil | $F_{3,151} = 2.52$<br>$P = 0.060$ | $F_{3,137} = 13.74$<br>$P = 2.1e-7$ | $F_{3,154} = 3.37$<br>$P = 0.040$ |
| | Subspecies x<br>Sterilized field soil | $F_{3,131} = 1.46$<br>$P = 0.23$ | $F_{3,118} = 2.72$<br>$P = 0.095$ | $F_{3,133} = 8.22$<br>$P = 0.00014$ |
| | Leaf number | $F_{1,462} = 1.47$<br>$P = 0.45$ | $F_{1,503} = 5.1$<br>$P = 0.073$ | $F_{1,486} = 1.12$<br>$P = 0.45$ |
| | Developmental Stage | $F_{1,474} = 5.71$<br>$P = 0.052$ | $F_{1,512} = 0.91$<br>$P = 0.34$ | $F_{1,493} = 4.49$<br>$P = 0.069$ |
| | Genotype [Subsp.] | $X^2_1 = 292.7$<br>$P = 3.9e-65$ | $X^2_1 = 37.05$<br>$P = 1.2e-9$ | $X^2_1 = 146.34$<br>$P = 2.2e-33$ |
| | Genotype x Sterilized<br>field soil | $X^2_1 = 0.46$<br>$P = 1$ | $X^2_1 = 0.03$<br>$P = 1$ | $X^2_1 = 0.79$<br>$P = 1$ |
| | Block | $X^2_1 = 9.28$<br>$P = 0.0047$ | $X^2_1 = 0.99$<br>$P = 0.32$ | |
| | Batch | | $X^2_1 = 3.52$<br>$P = 0.077$ | $X^2_1 = 4.28$<br>$P = 0.077$ |
| (b) |  | BC-ratio | Total [GS] | BC-diversity |
| | Genotype x<br>Soil microbiome | | $X^2_1 = 0$<br>$P = 0.97$ | |
| | Block | $X^2_1 = 3.33$<br>$P = 0.093$ | $X^2_1 = 3.96$<br>$P = 0.093$ | |
| | Batch | $X^2_1 = 0.01$<br>$P = 0.94$ | $X^2_1 = 6.74$<br>$P = 0.019$ | $X^2_1 = 11.18$<br>$P = 0.0025$ |

Both Total [GS] and BC-diversity varied among sterilized field soils, but not soil microbiomes (Figure 5a-b; Table 2). This indicates that the glucosinolate plasticity in response to whole natural field soils (Figure 4a) was primarily caused by variation in abiotic soil properties. Consistent with patterns of glucosinolate plasticity among natural habitats, the EAST subspecies was more responsive to abiotic soil variation than the WEST subspecies (Figure 5c-d; (Wagner and Mitchell-Olds, 2018a)). However, we detected no genetic variation for plasticity *within* subspecies in either the abiotic or biotic experiment (i.e., no genotype-by-treatment interactions; Table 2). Given that the same genotypes varied strongly in their plastic responses to environmental variation both within and among field sites (Wagner and Mitchell-Olds, 2018a), this suggests that glucosinolate plasticity in response to soil properties is less genetically variable than plasticity in response to other environmental factors that distinguish those field sites.

### Glucosinolate profiles of field-grown plants are partially congruent to those of greenhouse-grown plants

Our experiments demonstrated that variation among the soils of natural *B. stricta* habitats is sufficient to cause some plasticity of glucosinolate profiles. However, comparison to data from previously reported field experiments (Wagner *et al*., 2017; Wagner and Mitchell-Olds, 2018b) shows that other environmental factors also affect glucosinolate content of both leaves and roots. For each of the two greenhouse experiments, an independent field experiment using the same set of plant genotypes enabled us to test how the transition from field to greenhouse affected glucosinolate profiles.

In the “whole-soil” experiment, both root and leaf glucosinolate profiles differed between field-grown and greenhouse-grown plants, despite the use of field soil in the greenhouse (Figure 6a). In roots, BC-ratio was significantly lower in the greenhouse than in the field; however, this pattern was not observed in leaves. Notably, glucosinolate quantity was also affected. In roots, total concentrations of glucosinolates were substantially higher in the field than in the greenhouse, while the opposite pattern was observed in leaves. These results suggest that differences between field and greenhouse conditions–other than soil–induce organ-specific shifts in glucosinolate content.

**Figure 6:**
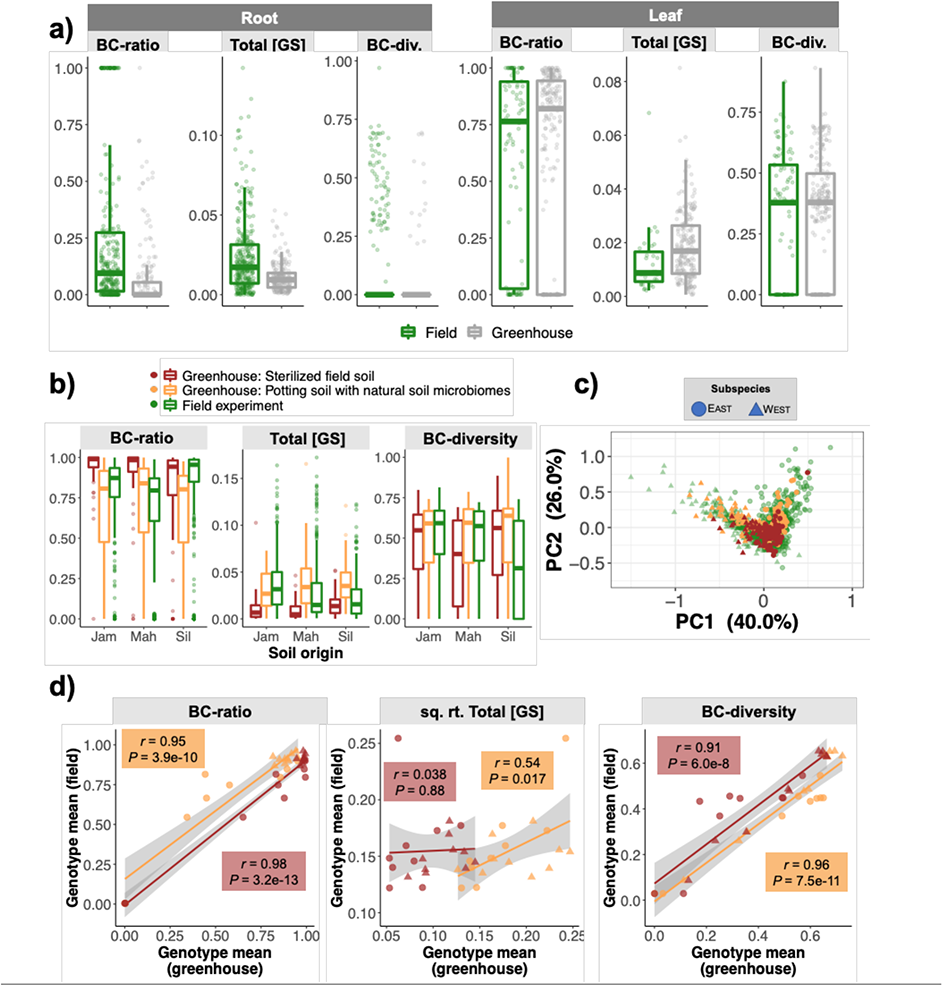
Comparison of glucosinolate profiles from plants grown under field conditions versus plants grown in the greenhouse in natural soils. Panel (a) shows root and leaf data from the “whole soil” experiment in which plants were grown in field-collected soils in the greenhouse. Ten high-[GS] outliers were omitted to improve plot readability. Panels (b-d) show leaf data from the “abiotic” and “biotic” experiments, in which plants were grown either in sterilized field soils or in sterilized potting substrate inoculated with microbial communities isolated from field soils. Five high-[GS] outliers were omitted for clarity. (c) Principal components analysis reveals that subspecies differences were the dominant factor explaining glucosinolate variation in all experiments. Colors denote experiments as in panel (b). Three high-PC2 outliers were omitted for the sake of plot clarity. (d) Genetic correlations between trait values measured in the greenhouse and field were weakest for Total [GS]. Colors denote experiments and shapes denote subspecies as in panels (b) and (c).

For the “biotic/abiotic” experiment, the difference in leaf glucosinolate phenotype between field and greenhouse conditions varied among traits and also depended on the habitat from which the soil treatments originated. For instance, BC-ratio was generally higher in plants grown in sterilized field soils, relative to field-grown plants and plants grown in potting substrate with natural soil biota; however, this pattern was observed for only two of the three field sites (Figure 6b). Similarly, we observed no difference in glucosinolate quantity between plants growing at Jackass Meadow and those grown in potting substrate inoculated with the soil biota from that field site; however, for the other two habitats, glucosinolate concentrations were considerably lower in field-grown plants. In contrast to the other glucosinolate traits, BC-diversity was largely unchanged between greenhouse and field conditions (Figure 6a-b).

Principal components analysis confirmed that genetic differences between the two subspecies are the strongest factor shaping overall glucosinolate profiles in all experiments (Figure 6c; see also (Novembre and Stephens, 2008)). Genetic correlations between trait values measured in the greenhouse and in the field were strong for BC-ratio and BC-diversity (Figure 6d), suggesting that most variation in these traits can be accurately predicted from the greenhouse regardless of which soil type is used.

However, the genetic correlation for Total [GS] measured in the field and the greenhouse was considerably weaker in the biotic experiment (*r*=0.54, *P=*0.017) and nonexistent for the abiotic experiment (*r*=0.038, *P=*0.88). This suggests that a sizable proportion of glucosinolate quantity variation cannot necessarily be extrapolated across environments, including soils. Together, these comparisons reflect the complexity of environmental variation that distinguishes these natural habitats and influences glucosinolate content in *B. stricta*.

### No evidence that sterilized field soils or soil microbial communities alter selection on glucosinolate profiles

Genotypic selection analysis did not support the hypothesis that natural soils or their microbial communities influence the adaptive value of any glucosinolate trait (*P* > 0.05 for all tests) under greenhouse conditions. In fact, we found little evidence for any relationship between relative fecundity and any of the three glucosinolate traits in either the biotic or abiotic experiment, with the sole exception of disruptive selection on BC-diversity observed in the biotic experiment (*P* = 0.027). These results contrast with observations from field experiments, in which Total [GS] was consistently under negative directional selection and BC-diversity was under either positive or negative directional selection—but not disruptive selection—in three of four habitats (Wagner and Mitchell-Olds, 2018a).

## DISCUSSION

### Abiotic causes of glucosinolate plasticity

Our findings demonstrate both the power and the limitations of reductionist approaches to identify specific environmental stimuli that affect plant phenotypes. Due to the complexity of natural habitats, we needed to manipulate individual environmental factors in a controlled setting in order to identify (or rule out) causes of glucosinolate plasticity that we had observed in the field (Anderson *et al*., 2014).

In this experiment, we found that purely abiotic soil variation is sufficient to cause plasticity of glucosinolate content. This is consistent with numerous observations that fertilization (especially with nitrogen and sulfur) can increase leaf glucosinolate content of Brassicaceous crops (Aires *et al*., 2006; Falk *et al*., 2007; Omirou *et al*., 2009; Kim and Juvik, 2011). For *B. stricta* growing in complex natural soils, however, the link between nutrient availability and glucosinolate quantity was not straightforward. For example, in both experiments, plants growing in Mahogany Valley soil had lower glucosinolate concentrations than plants growing in Silver Creek soil (Figures 4-5). This is true even though the Mahogany Valley soil contained higher levels of both nitrogen and sulfur, which are key nutrients needed for glucosinolate biosynthesis (Figure 2c). This suggests that the effect of any one nutrient on *B. stricta* glucosinolate profiles is likely contingent on levels of other nutrients available to the plant, as has been observed in *Brassica* crops (Omirou *et al*., 2009). More investigation into both the physiological limitations and the native habitats of *B. stricta* will be needed to understand how nutrient balance affects glucosinolate accumulation in this species, and how this relationship varies across the landscape.

Many other unmeasured soil variables, such as organic matter content and water potential, are also potentially important. Furthermore, the fact that greenhouse-grown plants differed from field-grown plants even when planted in field soils (Figure 6) suggests that aboveground abiotic variables such as temperature and UV radiation may shape glucosinolate content, as well. Analogous experiments could be conducted to confirm or rule out these other possible causes of glucosinolate plasticity.

### Soil microbiomes had no apparent effect on glucosinolate content

Our observation that soil microbes do not induce plasticity or selection of glucosinolate profiles may be somewhat specific to the relatively artificial experimental conditions (potting soil, lack of natural enemies, ample water, etc.). Microbes are likely to affect plant phenotype more strongly when conditions are more challenging (Bezemer and Vandam, 2005; Rolli *et al*., 2015). Nonetheless, these data do not support our initial hypothesis that natural variation in soil microbiomes *per se* would affect glucosinolate content. Our hypothesis was based on a body of evidence that root-associated microbes can directly regulate plant hormone signaling pathways, influence plant gene expression, and alter a wide array of ecologically important plant traits (Rashid and Chung, 2017). Previous studies found that a single bacterial strain can increase production of aliphatic glucosinolates via the jasmonate and ethylene signaling pathways in *Arabidopsis thaliana* (Brock *et al*., 2013; Pangesti *et al*., 2016). The results of single-strain experiments do not necessarily hold in more complex communities, however; it is possible that certain soil organisms may have had a similar effect on *B. stricta* physiology, while other community members had opposing effects, resulting in no net influence of the entire soil microbiome. On the other hand, we previously observed that soil microbiomes can speed or slow flowering time in *B. stricta*, indicating that these microbes do interact with the plant in some way, but with different implications for different phenotypic traits (Wagner *et al*., 2014). A more holistic survey of plant responses to soil microbiome variation—such as RNA-seq or metabolome profiling—would provide more information about the mechanisms by which *B. stricta* interacts with its belowground neighbors.

One important caveat is that by completely separating microbial communities from their natural substrates, this experiment may have missed interactions between soil organisms and soil chemistry that could have important implications for plant phenotype. For instance, imagining that the native Silver Creek microbiota can enhance phosphorus acquisition, this microbiome function may be detectable in the natural substrate (which is very low in phosphorus; Figure 2c) but not in the richer potting soil. Therefore, the lack of glucosinolate plasticity or selection in direct response to soil microbes in this experiment does not address whether microbes might affect glucosinolate profiles under other circumstances. A factorial experiment that combines several natural soils with several microbiomes might reveal different patterns.

### Glucosinolate content differs between greenhouse and natural settings

Finally, our results underline the importance of conducting experiments in natural conditions whenever possible. Despite our use of wild soils, the total glucosinolate concentration of our greenhouse-grown plants differed markedly from those of field-grown plants of the same genotypes (Figure 6). It should be noted that the field-grown plants used in these comparisons were actually germinated in potting soil before transplant to their natural habitats; therefore, these phenotypic differences probably result from some combination of soil and other variables such as light intensity, temperature, water availability, and non-microbial flora and fauna. It is likely that soil variation interacts with other environmental factors to shape plant phenotypes in complex natural habitats. This hypothesis could be tested by transplanting soils between field sites and measuring glucosinolate content in the field. Similarly, the lack of observed selection on glucosinolates in this experiment might reflect the absence of selective pressures in the greenhouse. Glucosinolates are important for defense against insects and pathogens (Hopkins *et al*., 2009; Pangesti *et al*., 2016; Stahl *et al*., 2016). They also are thought to contribute to drought resistance by regulating stomatal behavior (Khokon *et al*., 2011; Salehin *et al*., 2019) and by regulating aquaporin production in roots, thereby directly affecting water uptake and transport (Martínez-Ballesta *et al*., 2014). Our experimental plants received ample water and did not encounter natural predators.

Overall, our results show that soil variation is sufficient to cause plasticity of glucosinolate content in wild plant populations, which could ultimately affect the evolution of this important plant trait (Bradshaw, 1965; Wagner and Mitchell-Olds, 2018a). The ability for belowground pathogen and insect attacks to alter glucosinolate content of aboveground plant organs (and vice versa) is well appreciated (Bezemer and Vandam, 2005; Hopkins *et al*., 2009); however, direct effects of the soil on aboveground defenses should also be considered. For a more complete understanding of how glucosinolates shape biotic interactions in natural ecosystems, the reciprocal relationship—the effect of glucosinolate variation on plant-associated microbial communities—will be an interesting avenue of future research.

## DATA AVAILABILITY

All data and code used in this manuscript are freely available in a Zenodo repository (Wagner and Mitchell-Olds, 2022) accessible at https://doi.org/10.5281/zenodo.6998945.

## ACKNOWLEDGEMENTS

The authors thank Dr. Michael Reichelt for help identifying 6MTH and 4MOI3M, and Dr. Lauren Carley for helpful feedback on the manuscript. We thank J. Mays and the Duke University greenhouse staff for assistance with watering. MRW was supported by the National Science Foundation under Award Nos. IOS-2016351 and OIA-1656006 and by matching support from the State of Kansas through the Kansas Board of Regents. TMO was supported by grant no. R01 GM086496 from the National Institutes of Health.

